# Biological action at a distance: Correlated pattern formation in adjacent tessellation domains without communication

**DOI:** 10.1101/2021.07.14.452349

**Authors:** John M. Brooke, Sebastian S. James, Alejandro Jimenez-Rodriguez, Stuart P. Wilson

## Abstract

Tessellations emerge in many natural systems, and the constituent domains often contain regular patterns, raising the intriguing possibility that pattern formation within adjacent domains might be correlated by the geometry, without the direct exchange of information between parts comprising either domain. We confirm this paradoxical effect, by simulating pattern formation via reaction-diffusion in domains whose boundary shapes tessellate, and showing that correlations between adjacent patterns are strong compared to controls that self-organise in domains with equivalent sizes but unrelated shapes. The effect holds in systems with linear and non-linear diffusive terms, and for boundary shapes derived from regular and irregular tessellations. Based on the prediction that correlations between adjacent patterns should be bimodally distributed, we develop methods for testing whether a given set of domain boundaries constrained pattern formation within those domains. We then confirm such a prediction by analysing the development of ‘subbarrel’ patterns, which are thought to emerge via reaction-diffusion, and whose enclosing borders form a Voronoi tessellation on the surface of the rodent somatosensory cortex. In more general terms, this result demonstrates how causal links can be established between the dynamical processes through which biological patterns emerge and the constraints that shape them.

## Introduction

Central to current theories of biological organisation is a distinction between constraint and process. A constraint exerts a causal influence on a dynamical process and is not itself influenced by that process, at the spatial or temporal scale at which those dynamics take place. This definition permits a description of biological function in terms of constraint closure, i.e., the reciprocal interaction of constraints between processes operating at different timescales (***Montevil and Mossio, 2015; Montévil et al., 2016***; ***Mossio et al., 2016***) (see also ***Maturana and Varela, 1980***). A step towards falsifying such high-level descriptions of biological organisation is to formulate predictions at the level of specific biological systems, in which those predictions may be tested directly. To this end, our objective here is to operationalize the definition of constraint as causal influence on dynamical process.

The distinction between constraint and process is made explicit in the reaction-diffusion modelling framework (***Turing, 1952***), which has been successful in accounting for a wide range of biological (and other) phenomena, from the growth of teeth to the spread of tumors and the healing of skin (***Murray, 1984, 1989***). Reaction-diffusion models describe biological pattern formation in terms of local interactions amongst molecules or cells, which collectively amplify specific modes in an initially random distribution, with those modes determined by the relative size and shape of an enclosing boundary. Hence, the boundary shape is a constraint on the processes of short-range excitation and long-range inhibition from which pattern emerges.

Observing pattern contained by shape therefore suggests that the shape constrained pattern formation. But, alternatively, the enclosing shape may have emerged subsequently to, simultaneously with, or independently of, the formation of the pattern, and it is not obvious how to discriminate between these possibilities. One approach to establishing a causal influence of the boundary on the pattern is by *synthesis*. If the observed shape is imposed as a boundary condition for a reaction-diffusion model, and the evolution of that model gives rise to a similar pattern in simulation, we might infer a causal influence of the shape on the pattern. While compelling and important, such evidence is indirect, as computational modelling is limited to establishing existence proofs for the plausibility of hypotheses, rather than testing them directly. We seek therefore a complementary approach by *analysis* of the pattern, i.e., a direct means of testing between the hypothesis that the shape causally influenced the pattern and the null hypothesis.

To analyse an individual pattern in these terms, one could look for an alignment between the pattern and the boundary shape. For example, incrementing the diffusion constants from an initial choice that amplifies modes of the lowest spatial frequency will, on an elliptical domain, typically produce a sequence of patterns that is first aligned to the longer axis, and subsequently to the shorter axis. Indeed, for a well-defined boundary shape and a simple reaction-diffusion system generating a low spatial-frequency pattern, the alignment of an observed pattern to a hypothetical boundary constraint may be compared with a set of eigenfunctions derived from the linearized equations (i.e., using Mathieu functions for an elliptical domain; ***Abramowitz and Stegun, 1970***). But such methods break down for more complex boundary shapes, for higher-mode solutions, and for reaction-diffusion dynamics described by increasingly non-linear coupling terms.

In search of a more practical and robust method, the possibility we explore here is to exploit the fact that the shapes of adjacent biological domains are often related to one-another. That is, the processes that determine the shapes of adjacent domain boundaries may themselves be subject to common constraints, or indeed serve as constraints on one-another. Consider the following concrete example. In the plane tangential to the surface of the rodent cortex, the boundary shapes of large cellular aggregates called ‘barrels’ form a Voronoi tessellation across the primary somatosensory area (***Senft and Woolsey, 1991***). The barrel boundaries are apparent from birth, and from the eighth postnatal day develop ‘subbarrel’ patterns reflecting variations in thalamocortical innervation density (***Louderback et al., 2006***). A reaction-diffusion model, specifically the Keller-Segel formalism with its additional non-linear chemotaxis term, has been used to successfully recreate subbarrel structure in simulation, as well as to explain an observed relationship between the size of the enclosing barrel boundary and the characteristic mode of the subbarrel pattern (***Ermentrout et al. 2009***; see also ***Keller and Segel, 1971***). A synthetic approach has also helped establish that the barrel boundary shapes could emerge to form a Voronoi tessellation based on reaction-diffusion dynamics constrained by the action of orthogonal gene expression gradients on the processes by which thalamocortical axons compete for cortical territory (***James et al., 2020***). Hence in this system, the barrel boundary shapes that constrain subbarrel pattern formation via reaction-diffusion are thought also to be related by the common (genetic) constraints under which those barrel boundary shapes emerge.

Within such systems, the geometrical relationship between the shapes of adjacent domain boundaries might be expected to align the patterns that form within those boundaries to a degree that is reflected by the correlation between patterns in either domain. Hence measuring the degree of correlation between adjacent patterns could serve as a proxy for the degree of alignment to the boundary, and thus form the basis of a robust test for the hypothesis that shape constrained pattern formation.

Given chemical, mechanical, and other physical sources of spatial coupling in biological systems, it seems unlikely that pattern formation ever occurs completely independently in proximal and adjacent biological domains. But, in principle, how much of a relationship between patterns that form within adjacent domains might we expect to observe under the assumption that no communication occurs across domain boundaries?

On face value, this question might seem misguided. If pattern formation amongst cells within a particular domain occurs without the direct exchange of information with cells of an adjacent domain, then on what basis should we expect to measure any relationship at all between the patterns that form within adjacent domains? As we will show, strong correlations between patterns that self-organize independently in adjacent domains are in fact to be expected, if the shapes of those domains are geometrically related. That is, if the boundaries of adjacent domains abut, such that the domain shapes constitute a tessellation. Simulation experiments and analyses reported herein are designed to establish how relationships between domains on the basis of their shapes and common boundary lengths contribute to this somewhat paradoxical effect.

## Results

The key insight developed here is that patterns that self-organize independently in adjacent domains of a tessellation should nevertheless be correlated. Hence, by analysing the correlations between patterns measured in adjacent domains we can directly test the hypothesis that those observed patterns self-organized under constraints imposed by the observed borders within which they are enclosed. We will demonstrate the robustness of the (predicted) correlation effect via simulation, and show by analysis that it holds in a specific biological case (subbarrel patterning), but it is first instructive to present a toy example that reveals the effect most clearly (Fig. 1).

**Figure 1.**
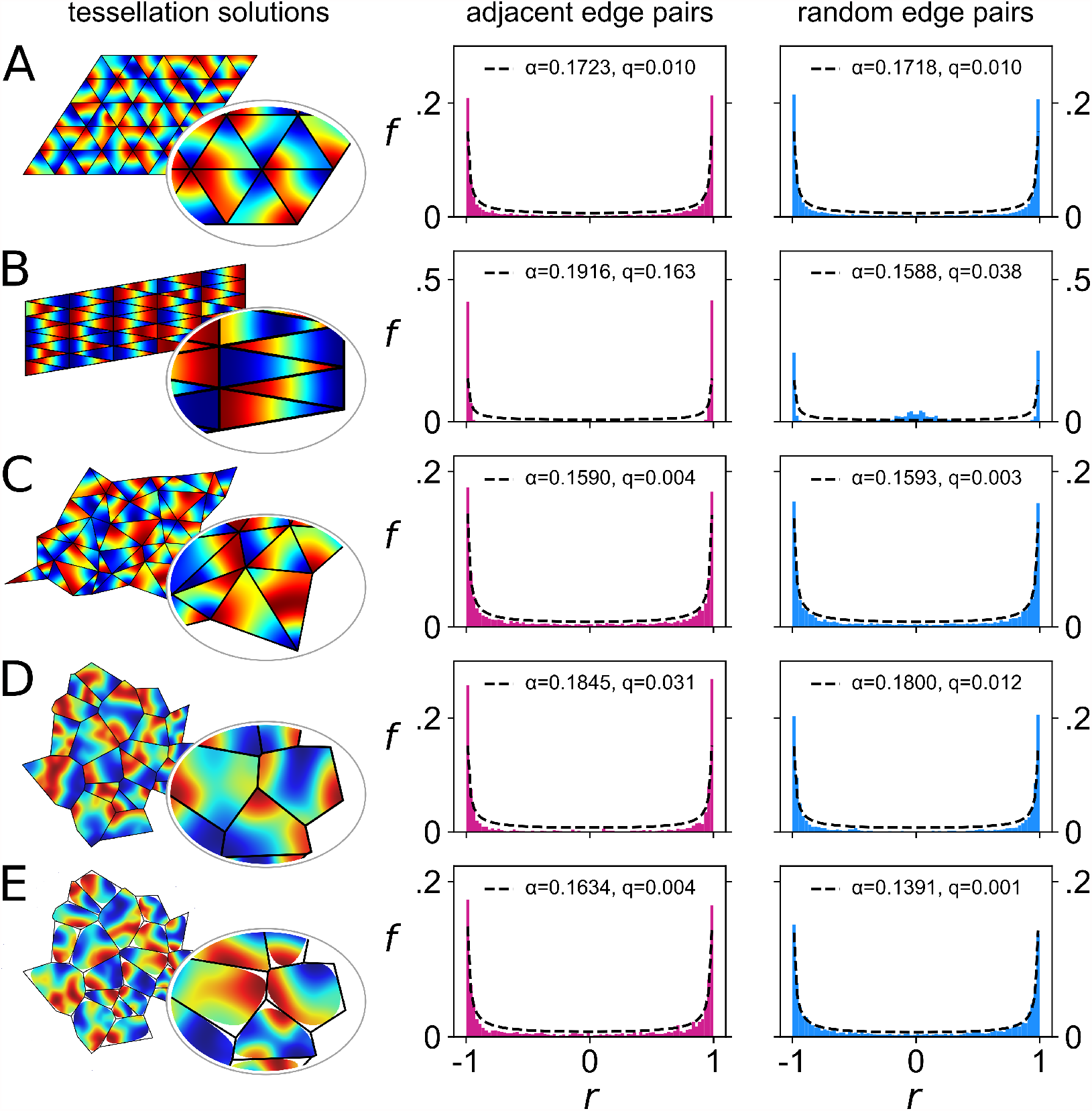
Correlated pattern formation in adjacent tessellation domains without communication. A system of reaction-diffusion equations (Eq. 2; *D*_*n*_ = 36, *χ* = 0) was solved using boundary shapes that tessellate in different ways (left column), with blue and red corresponding to extreme positive and negative values, and black lines delineating the domains. Values were sampled along the individual vertices of each domain and samples were correlated between edges of *different* domains, either amongst pairs of edges that are adjacent in the tessellation (center column) or randomly selected (right column). Histograms show the distributions (*f*) of correlation coefficients (*c*) obtained in either case, which were fit by the beta-distribution (dotted line) parameterized by *α* (see text for details; *q* is the sum of squared differences between the data and the fit). Rows **A**-**E** show data obtained from tessellations comprising domains with different shapes: **A** equilateral triangles; **B** isosceles triangles; **C** scalene triangles; **D** a Voronoi tessellation; **E** a Voronoi tessellation with rounded vertices. Peaks at ±1 in the histograms indicate that while pattern formation occurs entirely independently within each domain, patterns may become correlated between (adjacent) domains due to common constraints that derive from the fact that their boundary shapes tessellate.

### Bimodal pattern correlations amongst adjacent domains signal boundary constraints

Consider a reaction-diffusion system constrained by a boundary in the shape of an equilateral triangle (Fig. 1A). Solved for a choice of diffusion constants that yield patterns with the lowest spatial-frequencies, this system will generate one of two basic kinds of pattern. In the first, one of the extreme values of the reaction, positive or negative, will collect in one of the three corners of the triangle and values at the other extreme will be spread out across the opposite edge. Along that edge the values are essentially constant, and along the other two edges the values vary from extreme high to extreme low. In the second kind of pattern, values at the two extremes will collect in two corners and values around zero will collect in the third. Along one edge the values vary from extreme high to extreme low and along the other two they vary from zero to either extreme. Values sampled along the edges will vary between the two extremes in 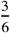 of the edge types (type 1), they will vary from one extreme to zero for 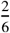 of the edge types (type 2), and they will not vary along the edge for 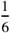 of the edge types (type 3). Assuming (for simplicity) that the two kinds of pattern occur equally often, and that pairs of edges are drawn at random from a large enough sample, two of the same edge type will be drawn with a probability of 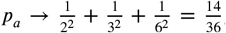, a type 1 and type 2 edge will be paired with a probability of 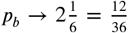, and a type 3 edge will be paired with a type 1 or 2 for the remaining 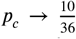. Now consider that along the edge, the magnitude of the correlation between the values sampled will be high for *p*_*a*_ pairs, low for *p*_*c*_ pairs, and intermediate for *p*_*b*_ pairs. Given that *p*_*a*_ *> p*_*b*_ *> p*_*c*_, that the magnitude of each correlation level increases with its probability of occurring, and that correlations and anti-correlations at each level are equiprobable given the symmetries within each kind of pattern, the distribution of correlations should be (overall) bimodal. Note that we describe the distribution as overall bimodal because smaller secondary peaks are expected to emerge around each distinct correlation level.

Consider next what happens when we substitute equilateral triangles with isosceles triangles (Fig. 1B). Reducing the number of axes of symmetry from three to one further constrains the kinds of patterns that are possible, causing (low spatial-frequency) solutions of the reaction-diffusion system to align with the perpendicular bisector of the base, and reducing the pattern along the edges to two types only. For example, if the base is the shorter side then 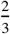 of the edges will be of type 1 and 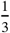 will be of type 3. Pairs of the same type constitute 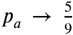 and pairs of different types constitute 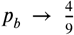, so again *p*_*a*_ *> p*_*b*_ and the distribution should again be (overall) bimodal. We note two important differences between the equilateral and isosceles cases. First, as pattern formation is more constrained by the isosceles boundary shape, and so the number of different kinds of patterns that are possible is reduced, the proportion of extreme correlations (and anti-correlations) has increased, from 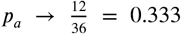 to 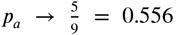. Second, the number of secondary peaks in the distribution of correlations has reduced to just two, around the positive and negative correlations corresponding to *p*_*b*_.

Now imagine laying the sample of equilateral triangles out, edge to edge and in no particular order, to form a regular tessellation. Clearly, when doing so, any triangle can be substituted or rotated so that a given edge is adjacent to any other, and hence we expect to sample from the same distribution of correlations whether we choose pairs at random, or limit our choices to those edges that are adjacent. But if we repeat the procedure for the isosceles triangles, this is no longer the case. Isosceles triangles only tessellate by arranging neighbours base-to-base or with the bases perpendicular bisectors antiparallel. A base cannot be adjacent to a non-base, and hence the distribution of correlations obtained from sampling adjacent pairs will lose its secondary peaks to display only the highest correlations and anti-correlations. So correlations sampled from adjacent rather than randomly selected edge pairs should be even more strongly bimodal.

Further, imagine randomly displacing each vertex of the tessellation of equilateral triangles in order to construct an irregular tessellation of scalene triangles (Fig. 1C). As each vertex is common to three triangles, each displacement changes the constraints on pattern formation in three triangles, from an initial minimally constraining configuration, and as such, increases the overall bimodality of the distribution of correlations. The irregular tessellation permits no substitution of domains, and hence, as in the isosceles case, we expect the overall bimodality of the distribution of correlations to be greater when comparing patterns amongst adjacent edges compared to randomly chosen edges.

An overall bimodal distribution of correlations amongst values sampled along pairs of edges from adjacent domains is therefore to be expected for domains that tessellate either regularly or irregularly. This property indicates that the domain boundaries constrained pattern formation. As a final thought experiment, consider that a jigsaw puzzle, i.e., an image into which borders are subsequently cut, will display perfectly strong positive correlations across adjacent edges and no anti-correlations. But our considerations thus far suggest that strong correlations and anti-correlations should be equally likely when the tessellation boundaries constrain subsequent pattern formation. Thus it is really the presence of strong anti-correlations in the distribution that evidences a causal influence of domain shape on pattern formation.

### Correlated pattern formation in adjacent domains of naturalistic tessellations

Considering pattern formation on tessellations of triangles is instructive, but to what extent do the considerations developed here apply to the kinds of tessellation observed in natural systems?

Examples of Voronoi tessellations are commonly found in the natural world (***Thompson, 1942; Honda, 1978, 1983***), including the packing of epithelial cells, the patterning of giraffe skins, and modular structures in the functional organization of the neocortex. The domains of a Voronoi tessellation enclose all points that are closer to a given ‘seed point’ than any other. As such, the polygonal structure of the tessellation is completely specified by a collection of seed points, with points along the polygonal boundaries equidistant to two seed points and points at the vertices equidistant from three. To test whether the predicted bimodal correlation is also to be expected in these naturally occurring tessellation structures, we generated random Voronoi tessellations from randomly chosen seed point coordinates, and solved the reaction-diffusion system (independently) within each domain. As shown in (Fig. 1D), the distribution of correlations sampled from along adjacent edges is again clearly overall bimodal. Hence, the effect is not specific to the case of triangles, and is to be expected for irregular tessellations of polygons that have a range of different numbers and arrangements of vertices.

The domains that comprise naturally occurring tessellations are often Dirichletform, but may not be strictly polygonal, with rounded corners rather than definite angles at the vertices (***Gómez-Gálvez et al., 2018***). And it is known that patterns formed by reaction-diffusion systems tend to be strongly influenced by the presence of definite angular intersections at the vertices (***Jung et al., 2017***). So to establish whether bimodality is also predicted for such natural structures, we reconstructed the random Voronoi tessellation and rounded the corners of the domains by joining the midpoints of each edge with quadratic Bezier curves whose first derivatives fit continuously at the midpoint. We then reconstructed the edges that corresponded to those of the original polygon by recording the points where the radial segments joining the centroid of the original polygon to its vertices cut the new shape. The reaction-diffusion system was solved again on the resulting domains, and (rounded) edges in the same locations as for the analysis of the original (polygonal) tessellation were correlated for a direct comparison. As shown in Fig. 1E, the distribution of correlations along adjacent edges is again predicted to be bimodal. Hence, the effect is not specific to polygonal domains and indeed is to be expected in this more general case.

### Emergence of bimodal correlations confirms that column boundaries constrain thalamocortical patterning in the developing barrel cortex

The emergence of subbarrel patterns in the rodent somatosensory cortex has been successfully modelled using the Keller-Segel reaction-diffusion system (***Keller and Segel, 1971***), with the borders of individual barrels imposed as a boundary constraint on pattern formation (***Ermentrout et al., 2009***). The barrel borders form a Voronoi tessellation, though the edges are typically a little rounded (***Senft and Woolsey, 1991***). The barrel structure is present from birth and the subbarrel patterns are first apparent at around postnatal day 8, and become clearly defined by around postnatal day 10, in stains for seretonin transporter and other markers for synaptic activity (***Louderback et al., 2006***). If subbarrel patterns emerge via reaction-diffusion dynamics under the constraints of the barrel boundaries, our analysis predicts that we should see a bimodal distribution of correlations along the common edges of adjacent barrels.

To test this hypothesis, we analysed images of seretonin transporter expression reported by Louderback and colleagues (***Louderback et al. 2006***; their Figure 4). The results of the analysis are shown in Fig. 2. We developed a simple computer program to sample the average image pixel intensity in rectangular bins pointing outward-normal to the two parallel sides of a user defined rectangle. Using this tool we defined rectangles to coincide with line segments corresponding to the septal regions that separate the barrels, then for each segment sampled from fifty bins along the outer edge of two adjacent barrel regions, to a depth of twenty pixels (∼ 85 *µ*m) into each of the barrels. Care was taken to ensure that the length of each line segment was as long as possible (to include as much of the border as possible), and that the width of each rectangle was as short as possible (to sample as close to the border of the barrels as possible), while not sampling from the septal region itself (to avoid introducing light/dark transitions into the sample that could cause spurious positive correlations). A small number of adjacent edges were excluded as their edges were not clearly parallel, but overall good coverage of the boundaries was achieved.

**Figure 2.**
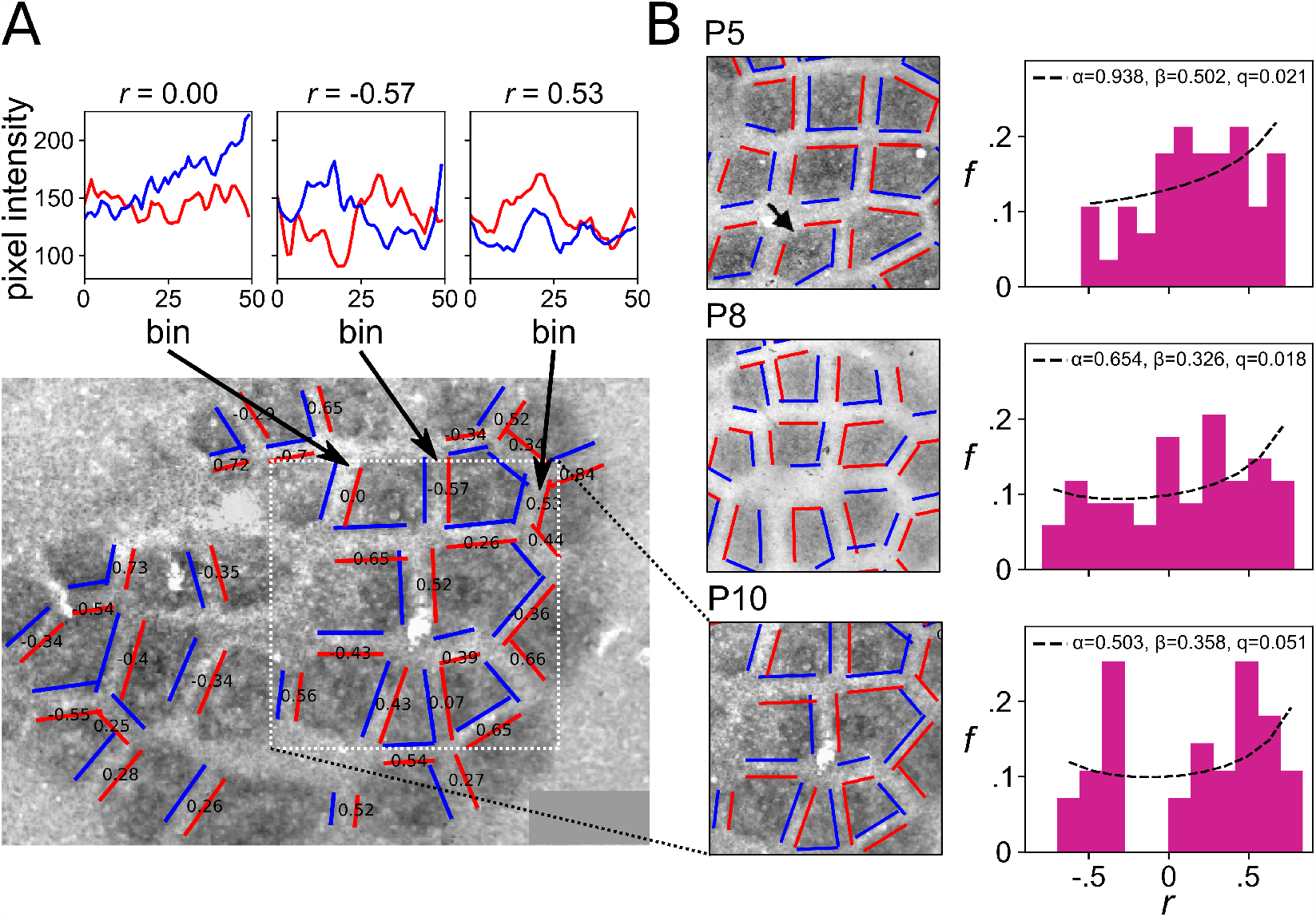
Emergence of correlated patterns in adjacent domains of the developing neocortex. We analysed images of immunohistochemical stains for serotonin transporter (5-HTT) expression on the surface of the rat barrel cortex, obtained at postnatal days 5 (P5), 8 (P8), and 10 (P10). This stain reveals the shapes of the barrel columns, each corresponding to a whisker on the animal’s snout, as large dark polygonal patches forming a Voronoi tessellation. From P8, sub-barrel structures become apparent and by P10 they clearly identify several regions of high synaptic density within many barrels. Panel **A** shows the details of the analysis method for the P10 image. Overlaid pairs of parallel red and blue lines show the extents along which image intensity was sampled for each pairwise comparison. Each line marks a vertex of the barrel boundary, and samples were constructed by averaging the grayscale intensity of pixels in one of 50 regularly spaced rectangular bins extending a short distance in from the line towards the corresponding barrel center. The correlation coefficient for each pair of samples is shown in black text, and the plots above show sampled data used for three example pairwise comparisons. Distributions of correlation coefficients obtained from pairs of edges from adjacent barrels are shown for each postnatal day in **B**, showing a clear progression from a unimodal shape at P5 to a bimodal shape at P10, and supporting the hypothesis that pattern formation within the barrels occurs postnatally and is constrained by the barrel boundary shapes. Original images from ***Louderback et al. (2006***).

Examples of the variation in pixel intensity along sampling bins spanning parallel line segments of adjacent barrels are shown at the top of Fig. 2A, revealing clear correlations and anticorrelations at postnatal day 10. In real data like this, it is conceivable that the technique could pick up spurious correlations, for example if image artefacts appeared in the sample from both edges of a pair, but we note that visible artefacts (e.g., circular bubbles of light or dark related to the underlying vasculature) very rarely spanned the width of the septa and when they did were very rarely located in or around the septa. Moreover, as noted above, anticorrelations are not to be expected by chance.

Images obtained from rats at postnatal days 5, 8, and 10 were analysed. At postnatal day 5, prior to when subbarrel patterns are reported to emerge, the distribution of adjacent-pair correlations in seretonin expression is unimodal, about a mean value of 0.18 ± 0.34. At postnatal day 8, when subbarrel patterns are reported to become apparent, two distinct peaks at a correlation of approximately ±0.5 are also apparent. At postnatal day 10, when subbarrel patterns are reported to be well defined, and are clearly visible in the image of seretonin expression, the distribution is clearly bimodal, with essentially all pairs showing non-zero correlations. Only the P10 distribution failed a test of unimodality (Hartigan’s dip test; *p* = 0.01). Thus our analysis supports the model of subbarrel pattern formation as a product of reaction-diffusion dynamics constrained by the barrel boundary shapes (***Ermentrout et al., 2009***). Moreover, this result demonstrates how the definition of constraint as a causal influence on biological process can practically be operationalized in terms of the distribution of adjacent pairwise pattern correlations, for reaction-diffusion systems on tessellated domains.

### Correlations are not bimodally distributed if borders are imposed after pattern formation

It was conceivable to us that a degree of correlation (and anti-correlation) may be expected due to chance for patterns of low spatial frequency, even without the boundary shape constraining pattern formation. To test for this we ran additional simulations, solving reaction-diffusion equations on an ensemble of regions from a randomly seeded Voronoi tessellation and applying no-flux boundary conditions on the edges of each region (Fig. 3A). We then solved the same system of equations on an ensemble of *circles*, centred at locations derived from the original Voronoi tessellation seed points, but subjected to additional random displacement by vectors whose radii and polar angles were chosen from a uniform distribution, normalised so that the new centres remained inside the original polygons. The size of each circle was chosen so that it minimally overlapped with the corresponding polygon from the original tessellation. We then overlaid the original tessellation onto the ensemble of circles, extracted the field values along the overlaid edges, and obtained the distribution of correlations for each case as previously described (Fig. 3B). The purpose of this procedure was to remove any possible influence of domain shape while ensuring that the data subjected to analysis were sampled from regions that tessellated precisely.

**Figure 3.**
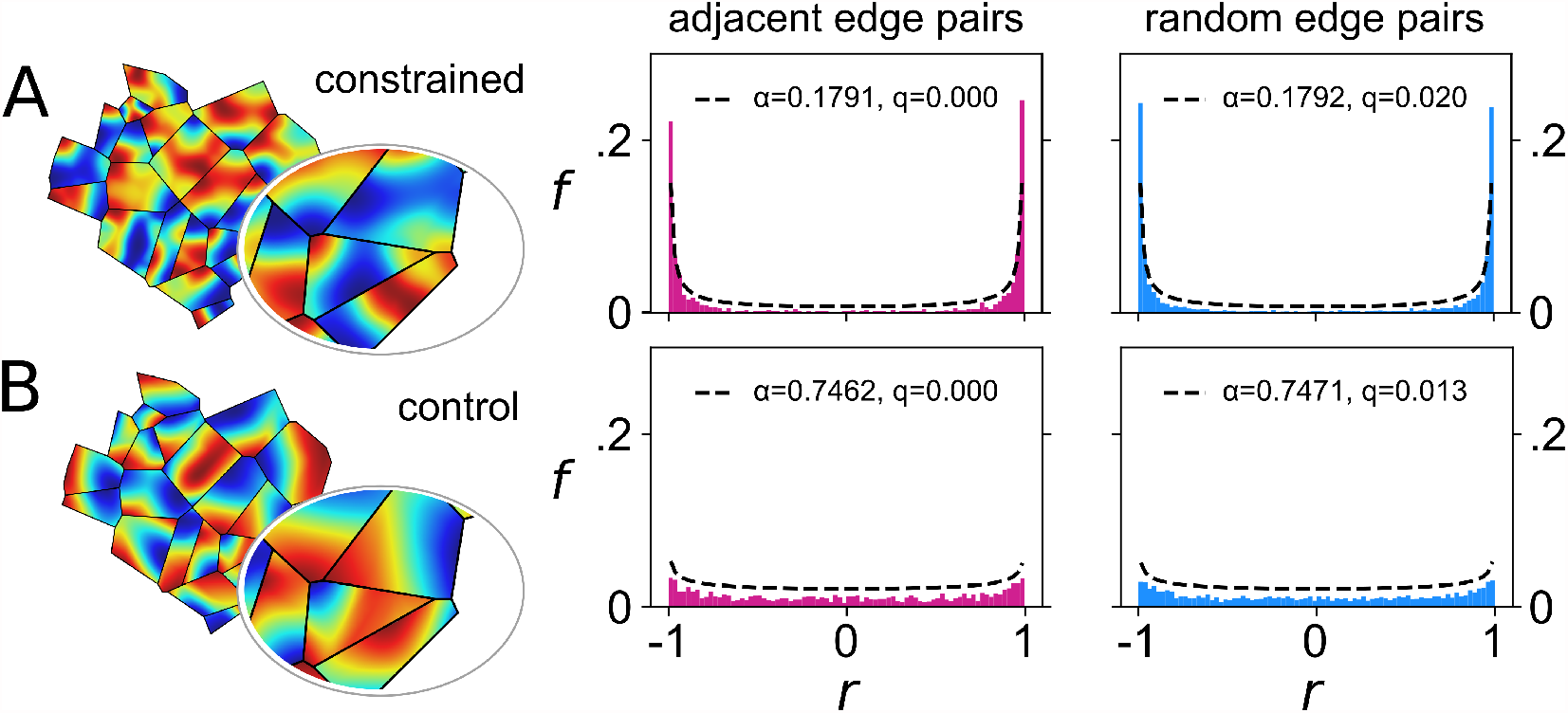
Analysis of control patterns formed without shaped boundary constraints registers only very weak correlations. Row **A** shows an analysis of patterns produced by the reaction-diffusion model (left column), based on correlations obtained for pairs of boundary vertices from adjacent (central column) or randomly matched (right column) domains of a Voronoi tessellation, in the case where the domain boundary shapes constrained pattern formation, as they do in Fig. 1. Row **B** shows a similar analysis in the case where the boundary shapes did not constrain pattern formation. Instead the same reaction-diffusion system was solved within circular domains containing no information about the tessellation, and the tessellation boundaries were imposed at the subsequent analysis stage only (regions of the circular boundaries that fall outside of the borders of the tessellation, shown as black lines, were discarded). Correlation distributions shown in **B** thus represent a control for the possibility that the bimodal distribution of correlations occurs by chance and/or is an artefact of the method of analysis. Only a very weak bimodality is present in this case, consistent with that to be expected based on a comparison with a model of random alignment that predicts a beta distribution with a high *α* coeffcient (see text for details). Pronounced bimodal peaks thus evidence the influence of the boundary shape as a constraint on pattern formation, as measured by a much lower *α* coeffcient in **A** (*α* = 0.179) compared to **B** (*α* = 0.747). Model parameters were *D*_*n*_ = 36 and *χ* = 0.

**Figure 4.**
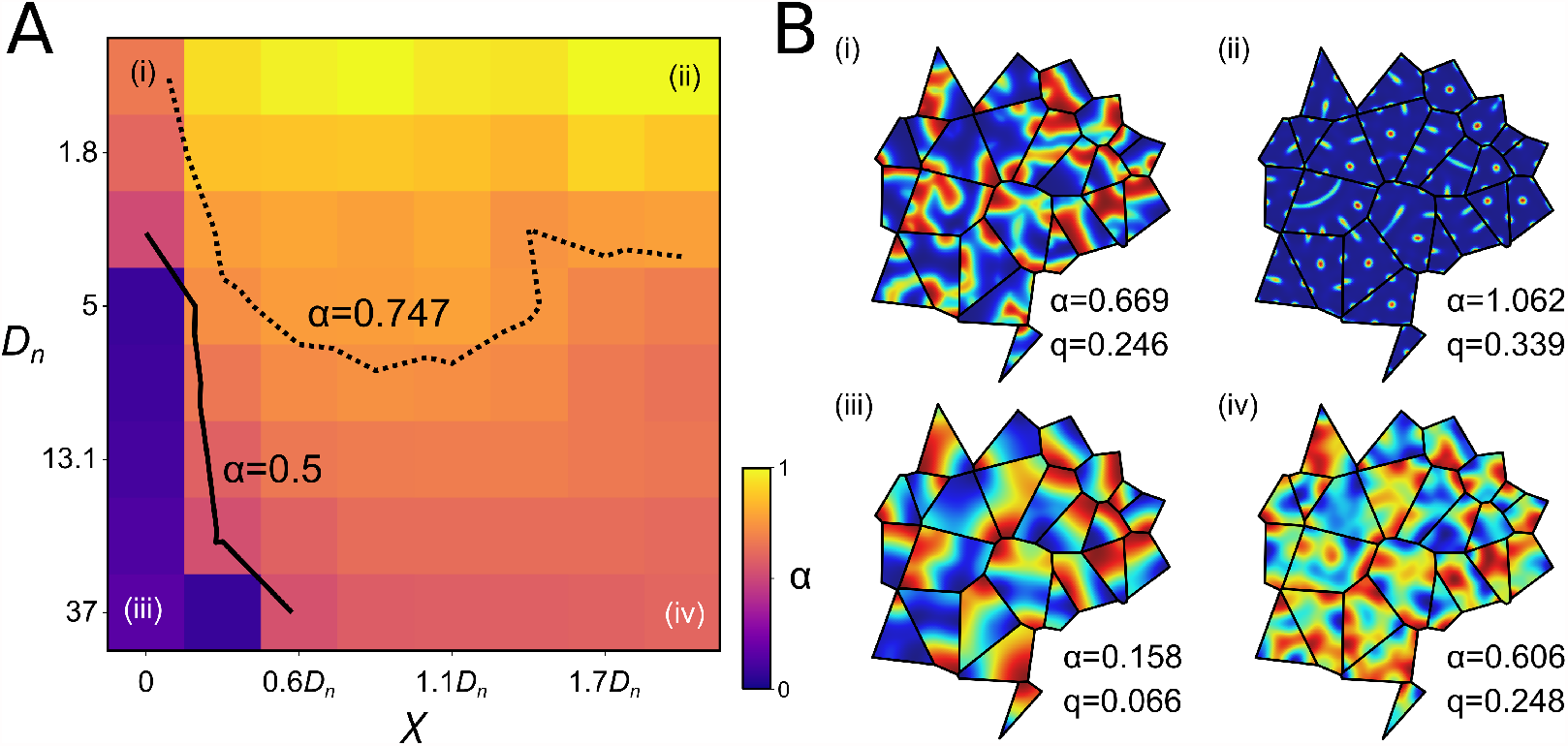
Correlated pattern formation in tessellated domains is predicted to emerge robustly across a wide range of pattern-forming systems. The magnitude of the correlations between adjacent domains induced by imposing Voronoi tessellation boundary shapes on pattern forming dynamics was measured in solutions obtained by using the Keller-Segel system across a wide range of parameters. Panel **A** shows values of *α* estimated from the distribution of 1000 pairwise correlation at each of sixty four combinations of parameter values. Combinations of eight values of the diffusion constant *D*_*n*_ and eight values of the constant *χ* that weights the non-linear coupling term were evaluated. The remaining free parameter, *D*_*c*_ was set to 0.3*D*_*n′*_ and increments in *χ* were expressed as proportions of *D*_*n*_ to cover a large parameter space. Note that when *χ* = 0, the system conforms to a classic reaction-diffusion system of the class originally studied by Turing. Overlaid contours correspond to thresholds in *α* below which the hypothesis that the domain boundaries constrained pattern formation should be accepted, based on the value obtained numerically from the control simulations reported in Fig. 3 (*α* = 0.747), or the overly conservative analytically derived value (*α* = 0.5). Based on the former, patterns are expected to be correlated by the tessellation boundary constraints across a large portion of the parameter space. Panel **B** shows example patterns for four extreme cases.

By visual inspection, the alignment between the patterns in Fig. 3B appeared similar to that formed under the constraints of the polygonal boundaries (Fig. 3A). However, histograms of the distribution of the correlations showed that they were quantitatively different. To understand this, we consider the known result (see e.g. ***Anderson 2009*** Ch. 4) that the scalar products of vectors that are uniformly randomly distributed on a unit hypersphere of dimension *D* − 1 (i.e. embedded in a space of dimension *D*) follow the beta distribution on *u* ∈ (0, 1),

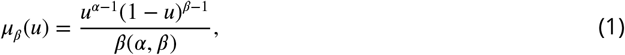

with *α* = *β* = (*D* − 1)/2, and *β* (*α, β*) the standard beta function (***Abramowitz and Stegun, 1970***). It can be seen that the beta distribution diverges at *u* = 0 and at *u* = 1 if *D* = 2, i.e. when the vectors are uniformly distributed on the unit circle, but that it conforms to a uniform distribution for a sphere in 3 dimensions (*D* = 3). Note that these dimensions pertain to the abstract vector space of all normalised edge vectors, and hence the dimension can in principle be as large as the numerical discretization that the tessellation permits. However, the coherence of the vectors derived from the smoothness of the solutions of Eq. 2 ensures that they lie in subspaces of much lower dimension. We measured correlations using the Pearson correlation coeffcient, which is equivalent to calculating the dot product of two unit vectors, and thus we can use simple algebra to map from the domain [−1, +1] to [0, 1]. If the edge vectors are not ‘pinned’ to the tessellation we expect them to be able to ‘slip’ relative to each other so they become uniformly distributed on a circle (see *Methods* for a proof). Estimates of the corresponding symmetric (*α* = *β*) beta distribution fits are shown with the histograms in Fig. 1 and Fig. 3A, where *α <* 0.5, from which we deduce that they are not uniformly distributed, exactly as expected if the influence of the tessellation on pattern formation were to preferentially select certain mutual orientations along adjacent edges. By contrast, the histogram obtained in the control condition (Fig. 3B) yields 0.5 *< α<* 1.0. Since the patterns that formed in this condition were not constrained by the tessellation, the increase in the degrees of freedom of their relative orientations produced a distribution that lost most of the bimodality and which thus approaches the uniform distribution. Note that replacing the coherent fields generated by reaction-diffusion with fields that have random values, and thus no spatial pattern, instead gives a distribution that approaches a normal distribution (*α* →∞).

The importance of this analysis is that it confirms that by estimating the value of *α* required to fit a beta distribution to a sample of edge correlations obtained experimentally, as in Fig. 2, we can deduce whether the boundary walls causally influenced the pattern formation. The value of *α* ∼ 0.75 obtained from the null model used to generate Fig. 3B serves as a practical threshold, below which such an influence is probable. While values below the analytically derived limit, *α <* 0.5, ascertain that the tessellating shapes served as constraints on pattern formation, the larger threshold value obtained numerically corresponds to a shift in *D* from 2 towards 3, which occurs in control simulations only when the domain centres are perturbed from the polygon centroids. Without such perturbations *α* → 0.5 (simulations not shown), and hence the analytical threshold of *α* = 0.5 may thus be considered overly conservative.

Applying this analysis to the data describing subbarrel pattern formation in Fig.2 at postnatal day 5 (P5) yields an estimate of *α* = 0.94 and *β* = 0.5, applying it at P8 yields *α* = 0.65 and *β* = 0.33, and at P10 the analysis yields *α* = 0.50 and *β* = 0.36. While the theory predicts that the distributions should be symmetric, i.e., *α* = *β*, the larger estimates for *α* reflect a general shift in each distribution to the right due to additional sources of positive correlation to be expected when extracting a fairly small sample from image data, as previously noted. As *α* is the parameter more sensitive to the presence of anticorrelations, we interpret its decrease from well above the numerical threshold at P5 to the analytical threshold at P10 as strong evidence that subbarrel patterns emerge postnatally under constraints imposed by the barrel boundary shapes.

### Alignment is robust to the form and spatial scale of the reaction-diffusion model

So far we have considered only the lowest mode solutions produced by a reaction-diffusion system. To explore whether the results should hold for the more complicated patterns that may be produced by more complex pattern-forming systems, we conducted a sweep of the parameter space, varying the diffusion parameter *D*_*n*_ and the parameter in Eq. 2 (see Methods) that weights the contribution of the non-linear coupling term, *χ*, while keeping *D*_*c*_ = 0.3*D*_*n*_ throughout (see Fig. 4). Note that when *χ* is set to 0, Eq. 2 specifies a classic Turing reaction-diffusion system, and that as *χ* increases the gradients induce a stronger nonlinear spatial effect, causing values greater than the field median to collect in small islands.

Fig. 4A shows the results of an eight by eight parameter sweep, where each point represents a unique combination of *D*_*n*_ and *χ*, and is coloured to show the estimated *α* values obtained by analysis of the resulting pattern correlations via Eq. 1. Contour lines corresponding to the analytic threshold (*α* = 0.5) and the empirical threshold (*α* ∼ 0.75) run approximately diagonally across the region, and the less conservative measure is effective at distinguishing the influence of the tessellation over more than half of this large parameter space. Example fields and the associated estimates of *α* are shown for the four extreme corners of the parameter space in Fig. 4B. Three are within the region where the empirical threshold can detect the effect of the tessellation on the solutions. In the top right, where *D*_*n*_ is low and *χ* is high, the fields become very concentrated and the nonlinear gradients in the region are so strong that the effects of the boundaries are not transmitted to the interior. However when either *D*_*n*_ is high or *χ* is low, parameters that yield complex fields that reflect the amplification of several modes clearly meet the criteria for accepting that the tessellation boundaries constrained pattern formation.

## Discussion

We have shown that because the shape of a domain boundary aligns pattern formation via reaction-diffusion, pattern formation within adjacent domains of a tessellation gives rise to an alignment between those patterns that can be measured as a strong (anti-)correlation between cells located on either side of a common boundary. Our simulation results demonstrate that the alignment of patterns in adjacent domains is predicted to be robust, with alignment occurring over a wide range of length scales, as set by the diffusion constants, and in reaction-diffusion systems both with and without non-linear coupling of the dynamic variables (Fig. 4). They also demonstrate that while rounding the vertices of the domains reduces the effect, it does not destroy it, and hence alignment is likely also to occur in biological domains where the boundary shapes may be less strictly polygonal (Fig. 1E). By showing that the effect is not to be expected in tessellated domains whose boundaries did not constrain pattern formation (Fig. 3B), our results establish bimodality in the distribution of correlations measured across adjacent edges of a tessellation (and specifically the presence of anti-correlations) as a useful test of the hypothesis that biological patterns formed via reaction-diffusion. This hypothesis was confirmed by the analysis of patterns of cell density that are thought to be formed via reaction-diffusion dynamics in the rodent somatosensory cortex, as a specific example system (Fig. 2).

The alignment effect is paradoxical, and an interesting biological example of action at a distance, because the process of pattern formation within a given domain occurs entirely independently of pattern formation in any other, and thus it involves no communication between cells that are located in different domains. Yet the effect is quite understandable, in geometric terms, when we consider that the boundary conditions of a given domain implicitly contain information about the boundary conditions of other domains, in the knowledge (or under the assumption) that those domains tessellate, and hence are related by a common underlying causal structure; e.g., by the collection of seed points from which a Voronoi tessellation originates.

The potential importance of this effect for understanding biological organization comes into focus when we consider how such causal structures might interact at different timescales (***Kauffman***, 1993; ***Rall***, 1996). Specifically, how might the alignment of patterns by their boundary constraints in turn constrain the slower processes that are involved in maintaining those boundary constraints? We can think of two broad answers, relating to the affordances of pattern alignment for material transport, and for information processing, though there may be several more.

In terms of material transport, if the pattern of concentration produced by a reaction-diffusion system corresponds to the density of cells or other physical obstacles, as it does in the example of neocortical patterning, then correlations along a common boundary edge create, in the regions of low concentration, channels through which other materials may flow. Uncorrelated patterns, such as those generated by our control simulations (Fig. 3B), are discontinuous at all borders and here create bottlenecks that restrict the flow of small materials and stop the flow of larger materials. In these terms, the anti-correlations that come with pattern alignment are of course bottle tops, permitting no flow at all, but with anti-correlations come correlations and thus the opportunity for unrestricted flow of small and large materials via the emergent channels. If transport of materials through these emergent channels participates in the maintenance of the objects that constitute the borders, for example by supplying them with energy or clearing their waste products, then the alignment of patterns by the boundary constraints in turn becomes a (useful) constraint on those boundary constraints.

As an interesting example, the Voronoi-like tessellation of dark patches that gives the giraffe skin its distinctive patterning is geometrically related to an underlying vasculature system. Particularly large arteries running between the patches supply a network of smaller arteries within the patches, which allow them to act as ‘thermal windows’ that efficiently radiate heat, and thus enable giraffes to thermoregulate in warm environments (***Mitchell and Skinner, 2004***). We note that giraffe panel substructures, not unlike subbarrel patterns in appearance, vary with the size of the panels, which in turn vary with the size of the animal in a manner predicted by reaction-diffusion modelling (***Murray, 1989***). This raises the intriguing possibility that a relationship between the structure of the vascular network and giraffe panel (and sub-panel) geometry may reflect a closure of constraints, co-opted for the thermal advantages it affords to these particularly large endotherms.

In terms of information processing, clustering of neurons to form tessellated patterns of cell density in and between brain nuclei constrains the transmission of signals between brain cells, and thus affords an opportunity for new information to be derived with reference to the underlying geometry, in turn enabling specific computations which facilitate survival (***Wilson and Bednar, 2015; Bednar and Wilson, 2016; Sterling and Laughlin, 2015***). The mammalian neocortex again provides a useful example. The arrangement of the barrels across the somatosensory cortex of rodents reflects the layout of the whiskers on their snouts, with cells of adjacent barrels responding most quickly and most strongly to deflection of adjacent whiskers. The relatively large size of the barrels, and the relatively slow velocities with which their efferents conduct action potentials, render downstream cells differentially sensitive to the relative timing of adjacent whisker deflections by virtue of their location with respect to barrel boundaries (***Wilson et al., 2011***). Neurons close to the borders respond selectively to coincident whisker deflections, and neurons that are closer to barrel A are selective for deflections of whisker B that precede deflections of whisker A by larger time intervals (***Shimegi et al., 2000***). As such, the system can use the underlying geometry to compute the relative time interval between adjacent whisker deflections via place-coding (***Jeffress, 1948; Izhikevich and Hoppensteadt, 2009***).

Within the additional cellular clusters that are formed via subbarrel patterning, neurons are tuned to a common direction of whisker movement (***Bruno et al., 2003***), and somatotopically aligned maps of whisker movement direction subsequently emerge, such that deflection of whisker A towards B selectively activates neurons of barrel A that are closest to barrel B (***Andermann and Moore, 2006***). This particular alignment of information-processing maps is thought to occur by the specific constraints that the barrel and sub-barrel geometry imposes on the otherwise general-purpose processes of reaction-diffusion and Hebbian learning by which cortical maps self-organize (***Wilson et al., 2010; Kremer et al., 2011***). The relationship between these two patterns that is suggested by the present results provides a potential geometrical basis by which downstream cells may report the coherence between information about patterns of movement through the whisker field derived from single whisker deflection directions and multi-whisker deflection intervals (***Kida et al., 2005***). The net effect is a representation of the ‘tactile scene’ that affords new possibilities for hunting and obstacle avoidance (***Jacob et al., 2008***).

There are many other examples of tessellated patterns in the brain, including spots and stripes in primate primary visual areas, and barrel-like structures in the brainstem, thalamus, and extrasensory cortical areas in rodents, as well as in various cortical areas in moles, dolphins, manatees, platypus, monkeys, humans, and more (see ***Manger et al. 1998*** for an overview). The precise role that these patterned modular structures (fields, stripes, barrels, blobs) might play in cortical information processing is yet to be fully characterised (***Purves et al., 1992; Wilson and Bednar, 2015; Bednar and Wilson, 2016***). However, in purely geometric terms, strong relationships between the shapes of cortical modules and the functional maps that they support have been well established. For example, iso-orientation contours radiating from the pinwheel centers that characterise topological maps of orientation preferences in primate primary visual cortex intersect with the boundaries of ocular dominance stripes at right angles (***Issa et al., 2008; Xu et al ., 2007***). And numerous features of these functional maps have been successfully modelled in terms of reaction-diffusion dynamics (e.g., see ***Swindale 1996***; ***Miikkulainen et al. 2005***; ***Wolf 2005***; ***Kaschube et al. 2010***). Hence, considering only neocortical patterning, it seems the opportunities for constraint closure in the brain via computational geometry are abundant.

***Montévil et al. (2016***) consider that reaction-diffusion dynamics introduce changes in the symmetries of biological systems that are *generic*, insofar as the nature of these changes derives from a restricted space of possibilities. As such, the changes in symmetry that are described by reaction-diffusion models are more akin to the well-defined phase transitions that occur in physical systems via symmetry-breaking, and they introduce a randomness in biological systems that is weak compared to the many other sources of randomness that shape biological organization. Generic constraints make the objects studied in physics suitable for mathematical analysis and synthesis (by computational modelling). However, for ***Montévil et al. (2016***), organized biological wholes are special objects that are, more importantly, also defined by constraints that are *specific*. That is, the dynamics of biological objects at any given moment, e.g., within an individual lifetime, depend on a history that spans ontogenetic and phylogenetic timescales in a way that precludes a meaningful *a priori* definition of their phase space, and thus precludes any meaningful analysis or synthesis (see also ***Kauffman, 2019, 2020***). Consequently, they suggest that modelling focused on deriving generic symmetries in biology will ultimately fail to capture the most important features, the individual accumulation of idiosyncrasies, that characterize biological wholes. As applied to reaction-diffusion modelling specifically, they warn that “We can describe a [symmetry change] explicitly with generic constraints but it is also possible to leave it implicit and consider that this single symmetry change is taken into account by the specificity of the object, among many other changes [… The] accurate description of any biological organism will always involve a component of specificity”.

We agree that this statement is accurate, and are sympathetic to this view of organisms, and other biological wholes, as historical objects (in context), but we do not agree with the suggestion that mathematical modelling in terms of such generic constraints is therefore fundamentally limited to describing biological parts and not biological wholes. The present results demonstrate how modelling at the level of generic symmetries/constraints can reveal meaningful insights at the level of biological wholes. Within the domains of a tessellation, pattern formation occurs by symmetry breaking on the timescale of the reaction-diffusion dynamics, but as patterns form a new symmetry emerges *between* the domains, and that new symmetry persists on the slower timescale at which the boundary conditions are defined and maintained. The alignment of patterns between domains constitutes a new (generic) symmetry insofar as it is invariant to the (specific) pattern that forms in either domain, as we have demonstrated most clearly using equilateral triangles. Thus the symmetry-breaking by which pattern forms in the shorter term gives rise to symmetries that persist in the longer term. As such, the opportunities that the alignment might afford to other processes (structural, transport, information-processing), persist at the same timescale at which the boundary conditions that gave rise to that alignment persist. If these opportunistic processes can help to maintain the boundary conditions, for example by channeling an external supply to the cells that form the boundary, then the cycle of constraints is closed, and a biological whole comes into being (c.f. ***Montevil and Mossio, 2015***).

Indeed, while “the best material model for a cat” may well be another cat, and preferably the same (specific) cat, we hope to have demonstrated here how the ‘imperfect, partial models’ advocated by ***Rosenblueth and Wiener (1945***), which like Turing’s reaction-diffusion model are targeted at understanding generic constraints on biological organization, remain useful tools for understanding biological wholes. Indeed, the alignment between reaction-diffusion processes in tessellated domains, and the possibility for constraint closure that this affords, may prove to be a useful theoretical model through which to explore, by analysis and synthesis, the fundamentals of biological organization.

## Materials and Methods

Patterns were formed on the two-dimensional plane **x** by solving reaction-diffusion equations of the form

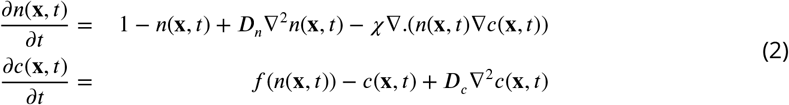

where *n* and *c* are two interacting species, *D*_*n*_ and *D*_*c*_ are diffusion constants, and the ‘chemotaxis’ term *χ* specifies the strength of the interaction between the two species. Following ***Ermentrout et al***. (***2009***) we use 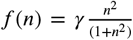 with *γ* = 5. For most of the simulations reported we set *χ* = 0, and thus implemented a standard reaction-diffusion system of the kind first studied by Turing, but to determine the robustness of the reported effects we also considered the more complex Keller-Segel equations, where *χ >* 0 (***Keller and Segel, 1971***).

Solutions were obtained numerically on a discretized hexagonal lattice of grid points using the finite volume method described by ***Lee et al. (2014***), and a Runge-Kutta solver was used to advance the solutions to a steady state (parameter values were chosen so that all solutions were eventually constant in time, i.e., patterns were stationary). Simulations were written in C++ with the help of the support library *morphologica* (***James and Wilson, 2021***) (see also ***James et al. (2020***)).

No-flux boundary conditions were applied at the edges of the regions derived from a given tessellation, by setting all the normal components of the spatial gradient terms in Eqs. 2 to zero. Importantly, the *tangential* gradients at the boundary were not constrained, allowing the patterns on either side of the boundary to represent the pattern in the whole domain while patterns across adjacent boundary walls had no constraints that might be correlated.

To compare the solutions along pairs of boundary vertices picked from two domains, the Pearson correlation coeffcient was calculated. Vectors **x**_1_ and **x**_2_ each contained the solution values in the hexagonal grid points along (and just inside) a boundary vertex from either domain, and these were combined to give the correlation coeffcient 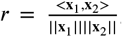, where the numerator represents the Euclidean scalar product and the denominator the product of the Euclidean norms. This operation can be thought of as measuring the angle between two unit normal vectors. As such, the distribution of the correlations may be considered a property of their distribution in a surrounding space – correlations will lie on a hypersphere whose dimension is between 1 and *D* − 1, with *D* the dimension of the containing space.

When comparing randomly matched edge vectors, the length of the shorter vector was increased to match that of the longer vector by linear interpolation. Results were verified using different resolutions of spatial discretization to ensure that the statistics were not influenced significantly by the coarseness of the discretization. A hexagon-to-hexagon distance of *O*(10^−3^) on a region whose spatial scale was normalized to be *O*(10^0^) was found to be sufficient. At this scale, regions contained *O*(10^2^) hexagons and edge vectors with length *O*(10^1^).

Using *α<* 0.5 as a threshold value for determining whether a set of patterns was constrained by the boundary shape can be justified analytically as follows. Consider a collection of edge patterns that are pinned to the vertices so that the maximal and minimal field values always appear at the two ends of each edge. The simplest (lowest mode) pattern that can be fitted to this constraint is a wave function cos(*θ*), where *θ* ranges from [0, *π*]. The correlation between two such functions is given by their dot product

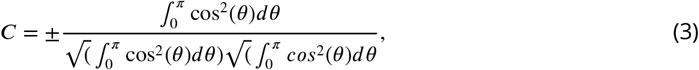

where the denominator gives the normalisation so that the vectors are of unit length, and hence Eq. 3 returns either *C* = 1 or *C* = −1. If we relax the constraint that the patterns must be pinned to the vertices and allow the pattern along each edge to shift by *ϕ* ∈ [−*π, π*], where *ϕ* is drawn from a uniform distribution, then we need to evaluate

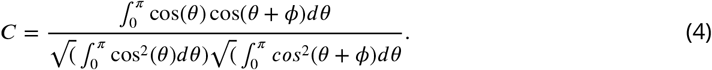

By the simple change in variables, *θ* = *θ* + *ϕ*, we can see that the denominator in Eq. 4 is the same as in Eq. 3. Expanding the numerator gives

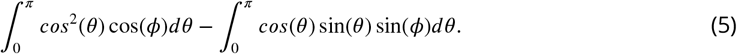

Either by explicitly evaluating the integral or by noting that cos(*θ*) and sin(*θ*) are anti-symmetric and symmetric about the midpoint of the range of integration, we can see that the second term vanishes, and hence that Eq. 4 becomes

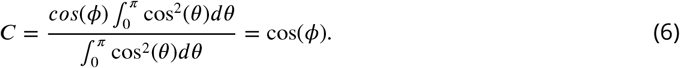

Therefore the distribution of correlations is cos(*ϕ*), where *ϕ* is a random variable drawn from a uniform distribution over [0, *π*] or over [*π*, 0], which is exactly equivalent to the distribution of the dot product between two vectors on the unit circle with an angle between them that is chosen from a random uniform distribution. Thus it gives a symmetric beta distribution with *α* = 0.5 (see e.g. ***Anderson*** (***2009***) Ch. 4).

Code for running the simulations reported in this paper is available at https://github.com/ABRG-Models/Tessellations.

## Acknowledgements

The authors thank Robert Schmidt at the University of Sheffield for useful discussion and comments on an earlier draft. This work was supported by a Collaborative Activity Award, Cortical Plasticity Within and Across Lifetimes, from the James S McDonnell Foundation (grant 220020516).

## Notes

https://github.com/ABRG-Models/Tessellations

### Competing Interest Statement

The authors have declared no competing interest.

